# Stat3 mediates Fyn kinase driven dopaminergic neurodegeneration and microglia activation

**DOI:** 10.1101/2024.07.05.602238

**Authors:** Sahiba Siddiqui, Fang Liu, Anumantha G. Kanthasamy, Maura McGrail

## Abstract

The Alzheimer’s Disease and Parkinson’s Disease risk locus Fyn kinase is implicated in neurodegeneration and inflammatory signaling. To investigate *in vivo* mechanisms of Fyn driven neurodegeneration, we built a zebrafish neural specific Gal4:UAS model of constitutively active FynY531F signaling. Using *in vivo* live imaging we demonstrate neural FynY531F expression lead to dopaminergic neuron loss and mitochondrial aggregation in 5 day larval brain. Dopaminergic loss coincided with microglia activation and induction of *tnfa*, *il1b*, and *il12a* inflammatory cytokine expression. Transcriptome analysis revealed Stat3 signaling as a potential Fyn target. Chemical inhibition experiments confirmed Fyn driven dopaminergic neuron loss and the inflammatory response were dependent upon activation of Stat3 and NF-κB pathways. Dual chemical inhibition demonstrated Stat3 acts synergistically with NF-κB in dopaminergic neuron degeneration. These results identify Stat3 as a novel downstream effector of Fyn signaling in neurodegeneration and inflammation.

**Summary Statement:** This study describes a novel *in vivo* model of neural Fyn Kinase activation and identifies Stat3 signaling as a downstream Fyn effector in dopaminergic neuron degeneration and neuroinflammation.

Graphical abstract

- Neural Fyn signaling drives dopaminergic neurodegeneration, mitochondria accumulation, and microglia activation
- Fyn driven neurodegeneration and cytokine expression are dependent on Stat3
- Stat3 and NF-kB pathways synergize in dopaminergic neuron degeneration

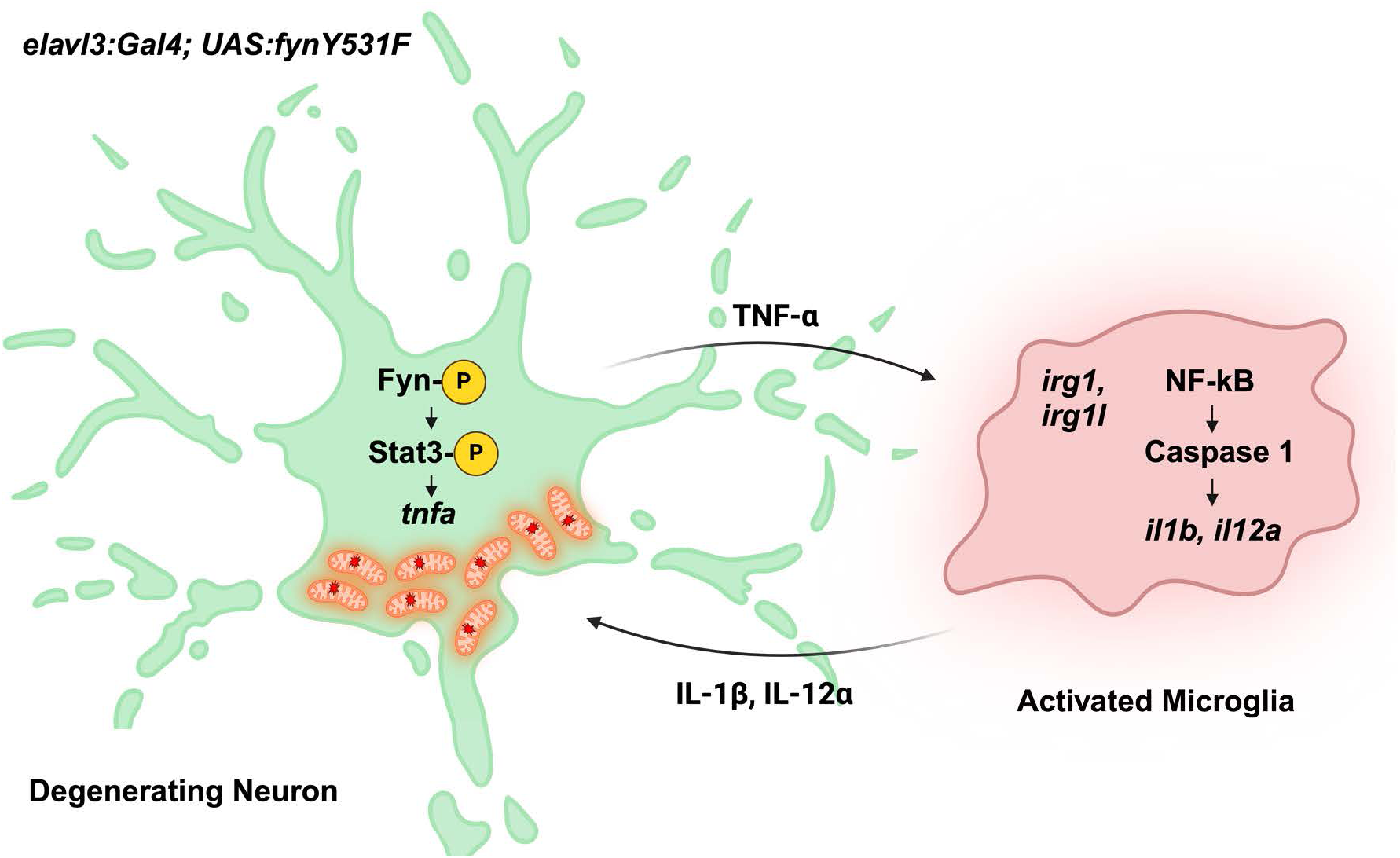

## Introduction

Cellular kinases play a central role in protein aggregation in dementia and neurodegeneration and their involvement in inflammatory signaling in Alzheimer’s Disease (AD) and related diseases (ADRD) such as Parkinson’s Disease (PD) is well established. The Src family member Fyn kinase has a key role in multiple neurodegenerative disorders (Guglietti et al. 2021). Fyn has been implicated in AD amyloid-β, tau and PD α-synuclein protein aggregate signaling (Nygaard 2018), and has been identified as a GWAS PD risk locus (Nalls et al. 2019). Increased Fyn expression and phosphorylation, a marker of Fyn activation, has been identified in AD and PD patient brain tissue (Low et al. 2021; Panicker et al. 2019; Guglietti et al. 2024), and Fyn activation has been reported to correlate with microglia activation (Panicker et al. 2019)*. in vitro* analyses of Fyn function indicate a role for Fyn signaling in both dopaminergic neurons and microglia activation in neurodegeneration. Fyn was shown to activate PKC-δ phosphorylation and oxidative stress induced cell death in rat dopaminergic neurons (Kaul et al. 2005; Saminathan et al. 2011). In microglia Fyn signaling stimulated inflammasome activation and cytokine induction through the PKC-δ/NF-κB pathway (Panicker et al. 2015; Panicker et al. 2019). In primary microglia Fyn leads to upregulation and phosphorylation of the Kv1.3 voltage-gated calcium channel, presenting a second pathway of Fyn driven microglia activation in neuroinflammation (Sarkar et al. 2020). Fyn knockout mice demonstrate a requirement for Fyn in neurotoxin induced dopaminergic neuron loss and microglia inflammatory response (Panicker et al. 2015). However, examining the mechanism by which elevated Fyn signaling drives neurodegeneration *in vivo* has not been fully investigated.

Zebrafish provides a powerful *in vivo* platform for modeling neurodegeneration (Chia et al. 2022). Neurotoxin and genetic zebrafish models of PD risk loci indicate conserved mechanisms underly dopaminergic neurodegeneration (Flinn et al. 2009; Flinn et al. 2013; Godoy et al. 2015; Kalyn et al. 2019; Omar, Kumar, and Teoh 2023). Dopaminergic neurons in larval and adult zebrafish forebrain are well described (Holzschuh et al. 2001; Wullimann and Rink 2001; Kaslin and Panula 2001) and A11-related dopaminergic neurons controlling mechano-sensation, locomotion and vision have been identified (Reinig, Driever, and Arrenberg 2017; Lohr, Ryu, and Driever 2009; Tay et al. 2011). Dopaminergic vDC clusters located in the posterior tuberculum of the ventral diencephalon until recently were considered analogous to the human DA neurons in the substantia nigra pars compacta (SNc) of the midbrain (Xi et al. 2011), which are lost in neurological diseases such as Parkinsons resulting in movement disorders. More recently, Th-positive neurons have been identified in the midbrain and in the hindbrain adjacent to the midbrain-hindbrain border, suggesting potential additional populations of dopaminergic neurons analogous to the human midbrain SNc controlling locomotion (Altburger, Holzhauser, and Driever 2023). Zebrafish transgenic reporter lines *dat:eGFP* (Xi et al. 2011) and *dat:mitoRFP* (Noble et al. 2015) for *in vivo* live imaging of dopaminergic neurons and mitochondria in the larval brain provide powerful tools for analysis of chemical and genetic models of dopaminergic neuron degeneration (Xi, Noble, and Ekker 2011; Godoy et al. 2015; Kalyn et al. 2019).

Using the binary Gal4; UAS system for cell type specific gene expression, we created an *in vivo* model of activated Fyn kinase signaling that drives neurodegeneration to investigate mechanisms of Fyn driven neurodegeneration. Neural specific expression of the constitutively active Fyn mutant Y531F led to larval morphological and phenotypic defects that recapitulate previously described zebrafish neurodegeneration models. Live imaging of larval brains revealed Fyn signaling drove loss of *dat:eGFP* labeled ventral diencephalon dopaminergic cell clusters and cell body aggregation of *dat:mitoRFP* labeled mitochondria over the course of 3 days to 5 days post fertilization. Dopaminergic neuron loss correlated with a shift in microglia morphology from ramified to amoeboid, consistent with elevated expression of the microglial activation marker *acod1*/*irg1*, and induction of inflammatory cytokine expression. Transcriptome analysis revealed alterations indicative of disrupted mitochondrial oxidative phosphorylation and metabolic pathways and revealed Stat3 as a novel Fyn downstream effector. Chemical inhibition reveals dopaminergic neuron loss and inflammatory gene expression were dependent on both Stat3 and NF-κB pathways. Together, Stat3 and NF-κB were shown to synergize with Fyn activated signaling in dopaminergic neuron loss. Our model of neural Fyn kinase signaling provides a new *in vivo* platform for investigating Stat3 targets and mechanisms of organelle stress in Fyn driven dopaminergic neurodegeneration.

## Results

### Zebrafish model of Fyn kinase driven neurodegeneration

To build a zebrafish model of Fyn-mediated neurodegeneration we used the two component Gal4-UAS system to drive overexpression of the constitutively active mutant FynY531F in neurons (Fig. 1). The zebrafish *fyna* Y531 residue is analogous to human Y528 located in the regulatory C-terminal domain of the protein. The Y528F phenylalanine phospho-null substitution mutation prevents the Fyn C-terminal tail from inhibiting the active site of the SH1 kinase domain, leading to a constitutively active kinase that drives persistent production of inflammatory cytokine Interleukin 1β (IL-1β) (Takeuchi et al. 1993). The zebrafish *fynaY531F* mutant cDNA was cloned into the *pTol2<14XUAS; gcry1:mRFP>* vector (Balciuniene et al. 2013) and two independent transgenic lines were established by standard Tol2 insertional transgenesis (Balciunas et al. 2006). The *fynaY531F* transgenic lines used in this study are *Tol2(14XUAS:fynaY531F; gcry1:mRFP)^is89^* (*UAS:fynY53F1is89*) and *Tol2(14XUAS:fynaY531F; gcry1:mRFP)^is90^* (*UAS:fynY531Fis90*).

**Figure 1.**
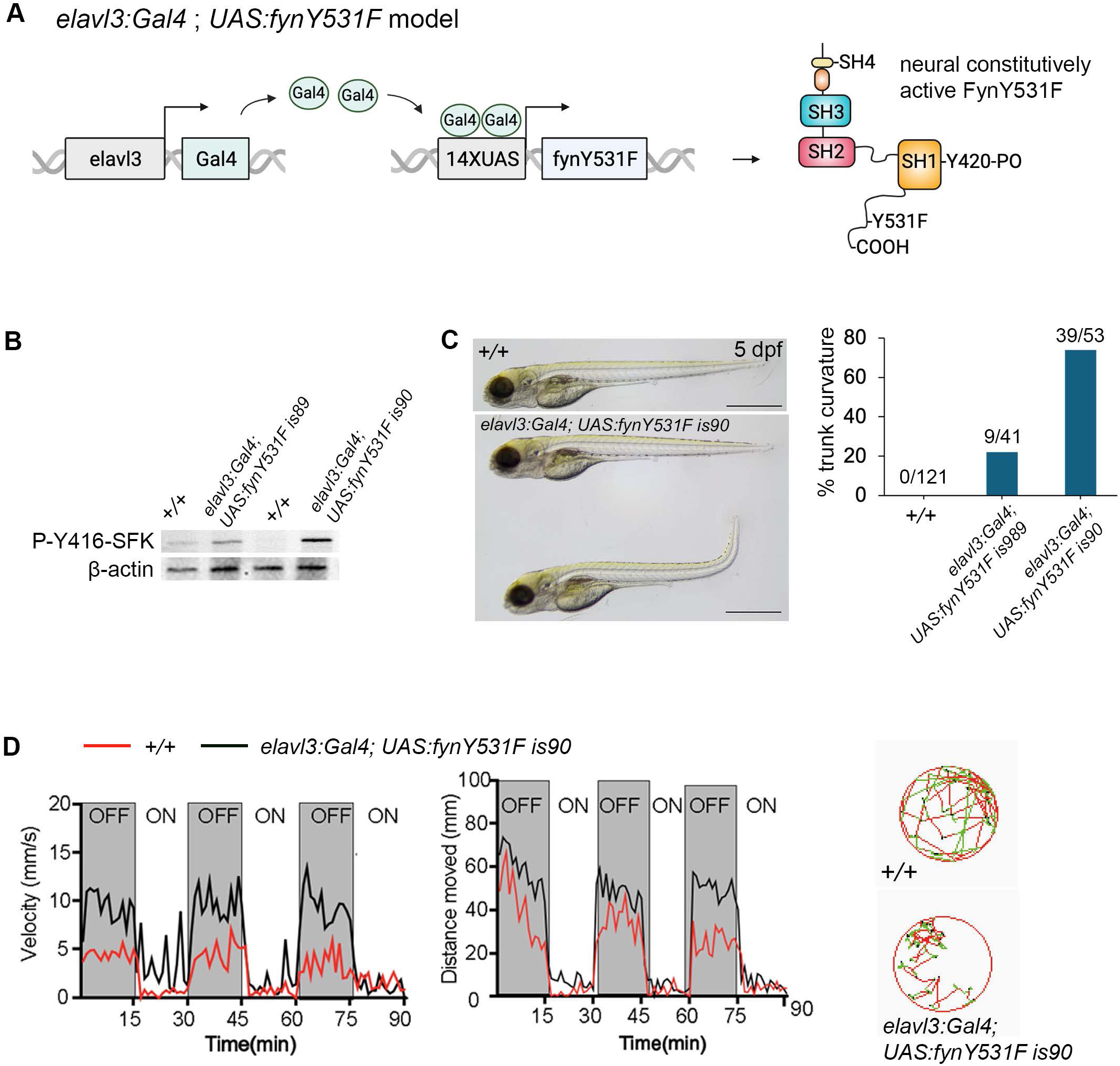
Gal4-UAS overexpression of FynY531F leads to autophosphorylation of Y416, trunk curvature, and swimming defects. **A** Diagram of *elavl3:Gal4; 14XUAS:fynY531F* neural specific model. **B** Western Blot probed with anti-P-Y416-SFK antibody showing elevated Y416 phosphorylation in *elavl3:Gal4; 14XUAS:fynY531Fis89* and *elavl3:Gal4; 14XUAS:fynY531Fis90* in comparison to wild type +/+ 5 dpf larvae. Anti-β-actin antibody was used as loading control. **C** Trunk curvature observed in a subset of *elavl3:Gal4; 14XUAS:fynY531Fis90*5 dpf larvae. Plot shows quantification of percentage of 5 dpf larvae showing trunk curvature in wild type +/+, *elavl3:Gal4; 14XUAS:fynY531Fis89*, *elavl3:Gal4; 14XUAS:fynY531Fis90*. **D** Measurement of velocity and distance moved in wild type +/+ and *elavl3:Gal4; 14XUAS:fynY531Fis90* 5 dpf larvae that showed normal trunk morphology. Scale bars 500 μm.

To determine if neural expression of constitutively active FynY531F would lead to neurodegeneration, we crossed the *UAS:fynY531F* lines with a neural specific Gal4 driver, Tg(*elavl3:Gal4=VP16)^nns6^* (Kimura, Satou, and Higashijima 2008) (designated *elavl3:Gal4*) (Fig. 1A). Western blots of extracts from *elavl3:Gal4;UAS:fynY531F* 5 day post fertilization (dpf) larvae were probed with an antibody that recognizes the autophosphorylation of tyrosine residue Y416 in the SH1 kinase domain of Src Family Kinases (p-Y416-SFK), corresponding to Y420 in Fyn kinase (Kouadir et al. 2012; Um et al. 2012; Wake et al. 2011). Neural specific expression of Y531F Fyn led to increased levels of p-Y416-SFK in comparison to wild type larvae (Fig. 1B), consistent with elevated Fyn kinase phosphorylation activity.

Neural expression of FynY531F resulted in 22% of *elavl3:Gal4; UAS:fynY5341Fis89* and 74% of *elavl3:Gal4; UAS:fynY531is90* 5 dpf larvae showing a curved trunk (Fig. 1C). To determine the impact of FynY531F overexpression on locomotory behavior, *elavl3:Gal4; UAS:fynY531Fis90* larvae with normal morphology were selected. The velocity and distance of light induced swimming over time was measured following previously established parameters (de Esch et al. 2012). Compared to wild type larvae, the *elavl3:Gal4; UAS:fynY531Fis90* larvae showed a reduction in both velocity and distance (Fig. 1D). These results indicate elevated levels of Fyn signaling in the *elavl3:Gal4; UAS:fynY531F* model correlate with morphological and motor defects that mimic previously described zebrafish models of neurodegeneration induced by the neurotoxin MPTP (1-methyl-4-phenyl-1,2,3,6-tetrahydropyridine) (Godoy et al. 2015; Kalyn et al. 2019; McKinley et al. 2005).

### Activated Fyn signaling leads to dopaminergic neuron loss and mitochondria accumulation

To determine whether the zebrafish neural FynY531F model led to neurodegeneration, live confocal imaging of dopaminergic neurons was performed in *elavl3:Gal4; UAS:fynY531F* larvae carrying the dopamine transporter (*dat*) GFP reporter *Tg(dat:eGFP)* (Xi et al. 2011) (Fig. 2). Compared to wild type *dat:eGFP* control larvae, at 3 dpf *dat:eGFP; elavl3:Gal4; UAS:fynY531F* larvae did not show a significant difference in the number of eGFP positive cells in the ventral diencephalic clusters (vDC) of dopaminergic neurons in the forebrain (Fig. 2A). In 5 dpf *dat:eGFP; elavl3:Gal4; UAS:fynY531F* larvae the level of dat:eGFP signal was reduced overall in the forebrain, midbrain, cerebellum and hindbrain, and the number of eGFP positive cells in the vDC clusters was significantly reduced compared to control (Fig. 2A). These results indicate activated FynY531F signaling leads to dopaminergic neuron loss, similar to the loss described in *pink* genetic (Flinn et al. 2013), *parkin* MO knockdown (Flinn et al. 2009), and MPTP neurotoxin (Godoy et al. 2015; Kalyn et al. 2019; McKinley et al. 2005; Xi et al. 2011) zebrafish models of neurodegeneration.

**Figure 2.**
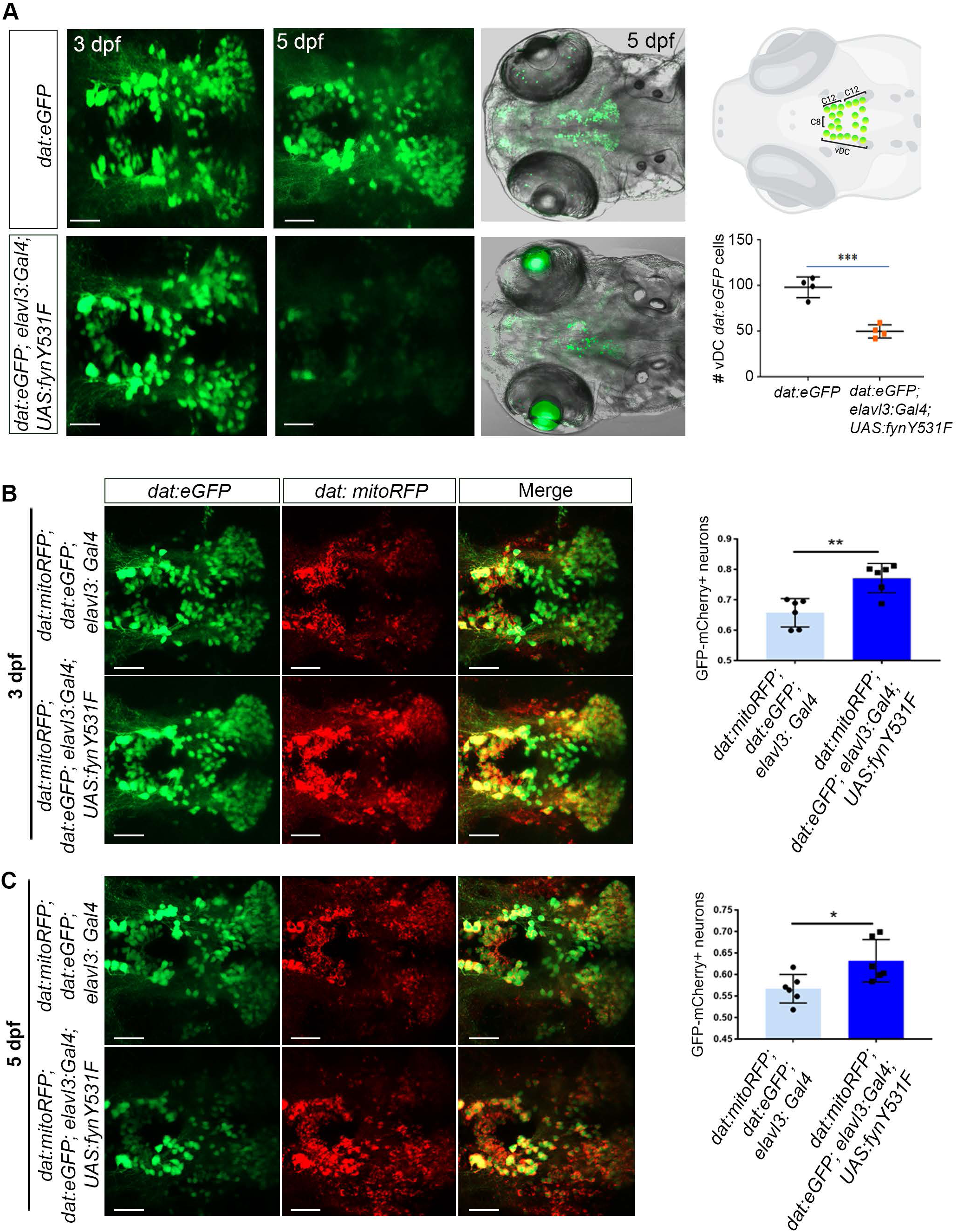
Zebrafish neural FynY531F overexpression drives dopaminergic neuron loss and mitochondria accumulation. **A** Live confocal imaging of ventral diencephalic neuron cluster (vDC) eGFP dopaminergic neurons in 3 dpf and 5 dpf control *dat:eGFP* and *elavl3:Gal4; UAS:fynY531F* 5 dpf larva. vDC in 5 dpf larval brain diagram adapted from Kalyn et al., 2019, created with BioRender.com. Quantification of vDC eGFP neuronal cell bodies in control *dat:eGFP* and *dat:eGFP; elavl3:Gal4; UAS:fynY531F* 5 dpf larva (n=4). **B** Live confocal imaging of eGFP and mCherry in vDC neurons in control *dat:mitoRFP*; *dat:eGFP*; *elavl3:Gal4* and *dat:mitoRFP*; *dat:eGFP*; *elavl3:Gal4; UAS:FynY531F* 3 dpf larval brain. Quantification of overlap of eGFP and mCherry signal in vDC neuron cell bodies in control *dat:mitoRFP*; *dat:eGFP*; *elavl3:Gal4* and *dat:mitoRFP*; *dat:eGFP*; *elavl3:Gal4; UAS:FynY531F* 3 dpf larva (n = 6). **C** Live confocal imaging of eGFP and mCherry in vDC neurons in control *dat:mitoRFP*; *dat:eGFP*; *elavl3:Gal4* and *dat:mitoRFP*; *dat:eGFP*; *elavl3:Gal4; UAS:FynY531F* 5 dpf larval brain. Quantification of overlap of eGFP and mCherry signal in vDC neuron cell bodies in control *dat:mitoRFP*; *dat:eGFP*; *elavl3:Gal4* and *dat:mitoRFP*; *dat:eGFP*; *elavl3:Gal4; UAS:FynY531F* 5 dpf larva (n = 6). Statistical analysis was performed with two-tailed unpaired Students t-test. Bars represent mean +/- SEM. * p<0.05; ** p<0.01. Scale bars 100μm.

To examine whether activated Fyn signaling impacts mitochondria in dopaminergic neurons, live confocal imaging of the dopamine mitochondrial fluorescent mCherry reporter line *Tg(dat:tom20 MLS-mCherry) (dat:mitoRFP)* (Noble et al. 2015) was performed in control *elavl3:Gal4* and *elavl3:Gal4; UAS:fynY531F* larvae at 3 dpf (Fig. 2B) and 5 dpf (Fig. 2C). 3 dpf *dat:mitoRFP; elavl3:Gal4; UAS:fynY531F* larvae showed an increase in mCherry signal co-localizing with eGFP in dopaminergic vDC cell bodies compared to control larvae (Fig. 2B; p<0.002). At 5 dpf colocalization of mCherry signal with eGFP in remaining vDC cell bodies was significantly higher than in controls (Fig. 2C; p<0.03), despite the reduction in overall vDC number in *dat:mitoRFP; elavl3:Gal4; UAS:fynY531F* larvae. The increase in *dat:mitoRFP* signal suggests mitochondria accumulate in the dopaminergic cell bodies of *elavl3:Gal4; UAS:fynY531F* larvae. This indicates elevated Fyn signaling leads to neuronal degeneration due to a defect in mitochondrial turnover or dysfunction.

### Activated Fyn leads to induction of inflammatory cytokines and microglia activation

To test whether FynY531F driven dopaminergic neuron loss in the larval brain correlated with microglia activation, a marker of neuroinflammation, 5 dpf control +/+ and *elavl3:Gal4; UAS:fynY531F* larvae were labeled with an anti-4C4 hybridoma supernatant (Fig. 3A). The 4C4 hybridoma supernatant recognizes galectin 3 binding protein and labels a subset of brain microglia (Mazzolini et al. 2020; Becker and Becker 2001; Chia et al. 2018; Rovira et al. 2023). Compared to wild type controls, the forebrain and midbrain of *elavl3:Gal4; UAS:fynY531F* larvae did not show a significant difference in the number of 4C4-positive microglia in the forebrain (p=0.39) and midbrain (p=0.12), while the number detected in the hindbrain was elevated (p<0.0001) (Fig. 3A, B). However, in comparison to the ramified appearance in wildtype +/+ controls, the 4C4 positive microglia in *elavl3:Gal4; UAS:fynY531F* larval brains appeared amoeboid in morphology (Fig. 3C). The change in microglia morphology from ramified to amoeboid suggests microglia become activated in response to elevated Fyn signaling in the larval brain.

**Figure 3.**
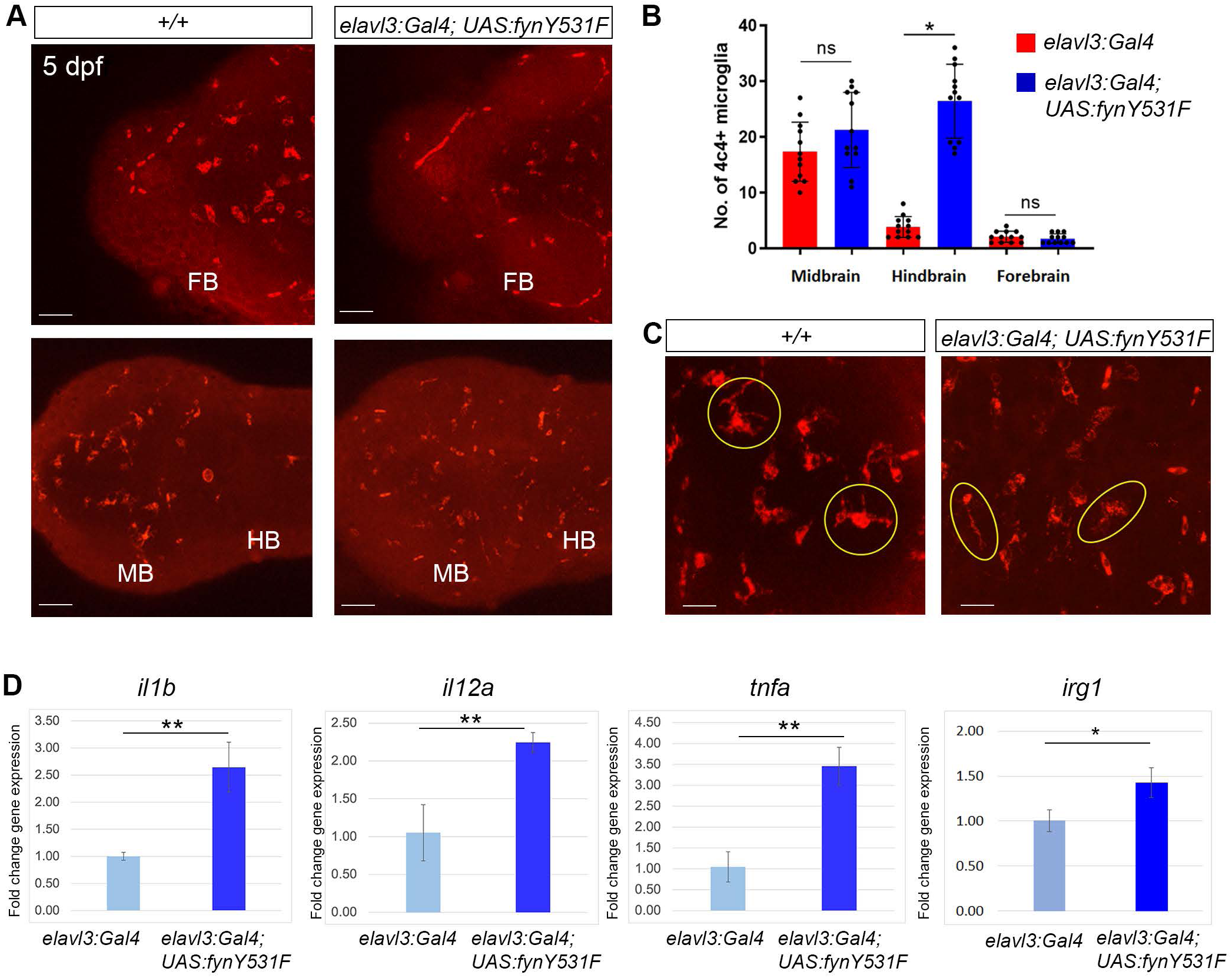
Zebrafish neural FynY531F overexpression drives microglia activation and cytokine induction. **A** 4C4 immunolabeling of microglia in wild type +/+ and *elavl3:Gal4; UAS:fynY531F* forebrain and mid/hindbrain. **B** Quantification of 4C4 labeled microglia in *+/+* and *elavl3:Gal4; UAS:fynY531F* 5 dpf larva brain (n=12). **C** Higher magnification images of *+/+* ramified and *elavl3:Gal4; UAS:fynY531F* amoeboid 4C4-labelled microglia. **D** RT-qPCR of *il1b*, *il12a*, *tnfa*, and *irg1* on RNA extracts from control *elavl3:Gal4* and *elavl3:Gal4; UAS:fynY531F* 5 dpf larvae (n= 3 biological replicates for each genotype). Statistical analyses were performed with two-tailed unpaired Students t-test. Bars represent mean +/- SEM. ns not significant; * p< 0.05; ** p< 0.01. Scale bars A 50μm; C 20μm.

To determine whether activated Fyn signaling led to induction of gene expression indicative of inflammatory signaling and microglia activation, RT-qPCR was used to examine the expression level of the cytokine genes IL-1β (*il1b*), IL-12α (*il12a*) and TNF-α (*tnfa*) in dissected head tissue of 5 dpf control *elavl3:Gal4* and *elavl3:Gal4; UAS:fynY531F* larvae. *elavl3:Gal4; UAS:fyn531F* larvae showed a significant increase in the levels of *il1b* (p=0.003), *il12a* (p=0.006), and *tnfa* (p<0.002) (Fig. 3D). The gene encoding the activated microglia/inflammatory macrophage marker Aconitate decarboxylase 1/Immuno-Responsive Gene 1 (*acod1/irg1*) also showed significant elevation (p<0.02) in *elavl3:Gal4; UAS:fynY531F* larvae (Fig. 3D). Together these results suggest elevated neural Fyn signaling driving dopaminergic neuron loss correlates with induction of inflammatory cytokine expression and microglia activation.

### *elavl3:Gal4; UAS:fynY531F* dopaminergic neuron loss and microglia activation are dependent on Fyn kinase activity

In order to demonstrate dopaminergic neuron loss, cytokine elevation, and microglia activation are dependent on constitutive Fyn kinase signaling in the *elavl3:Gal4; UAS:fynY531F* model, we tested whether the Src family kinase inhibitor Saracatinib could suppress these cellular and molecular phenotypes (Figure 4). 3 dpf *dat:eGFP* control and *dat:eGFP; elavl3:Gal4; UAS:fynY531F* larvae were treated continuously with 20uM Saracatinib for 2 days and collected at 5 dpf for live confocal imaging of vDC dopaminergic neurons and RNA extraction for RT-qPCR (Fig. 4A). Live imaging of eGFP expression in 5 dpf Saracatinib treated larvae showed suppression of vDC neuron loss (Fig. 4B) and retention of vDC neuron numbers at levels equal to control mock (p=0.15) or Saracatinib-treated *dat:eGFP* (p=0.94) larvae (Fig 4C). Western blot analysis of Saracatinib-treated 5 dpf larvae showed a reduction in the level of P-Y416-SFK in comparison to DMSO mock treated controls (Fig. 4D), indicating FynY531F kinase activity was suppressed. RT-qPCR revealed Saracatinib treatment led to a reduction in *il1b* and *il12a* expression in FynY531F *elavl3:Gal4; UAS:fynY531F* larvae, demonstrating elevated inflammatory cytokine levels were dependent on activated FynY531F signaling (Fig. 4E). In contrast, neither elevated *tnfa* nor *irg1* levels were suppressed by Saracatinib treatment of *elavl3:Gal4; UAS:fynY531F* larvae (Fig. 4E), possibly due to the non-specific action of the general Src Family kinase inhibitor causing an impact on *tnfa* nor *irg1* levels.

**Figure 4.**
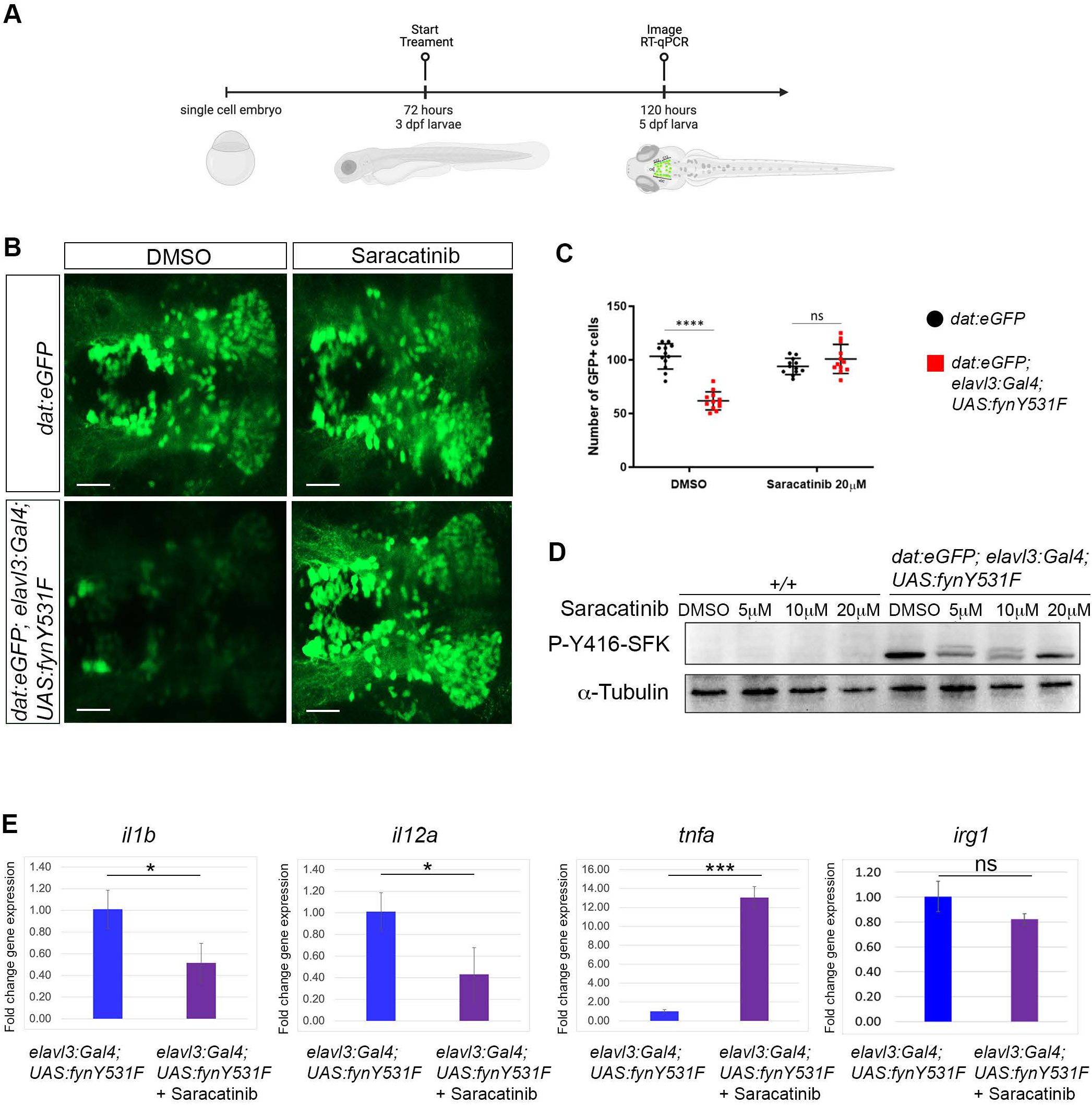
Src family kinase inhibitor Saracatinib suppresses FynY531F driven dopaminergic neuron loss, Fyn activation, and *il1b*, *il12a* cytokine induction. **A** Time course of Saracatinib treatment beginning at 72 hpf. **B** Live confocal imaging of vDC cluster eGFP dopaminergic neurons in control *dat:eGFP* and *elavl3:Gal4; UAS:fynY531F* 5 dpf larva mock treated with DMSO or treated with 20mM Saracainib. **C** Quantification of vDC eGFP positive neuronal cell bodies in control *dat:eGFP* and *dat:eGFP; elavl3:Gal4; UAS:fynY531F* 5 dpf larva mock treated with DMSO or treated with 20mM Saracatinib (n=12). Analysis was performed using two way ANOVA with Tukey’s multiple comparison. **D** Western blot of 5 dpf larval extract from wild type and treated with DMSO and increasing amounts of Saracatinib, probed with anti-Srk family kinase Y416PO. Anti acetylated tubuling and anti-alpha-Tubulin were used as loading controls. **E** RT-qPCR of *il1b*, *il12a*, *tnfa*, and *irg1* on RNA extracts from untreated and 20μM Saracatinib treated 5 dpf *elavl3:Gal4; UAS:fynY531F* 5 dpf larvae (n= 3 biological replicates all genotypes and conditions). RT-qPCR was analysis was performed with Two-tailed unpaired Student’s t-test. Bars represent mean +/- SEM. ns, not significant; * p< 0.05; *** p< 0.001. Scale bars 100μm.

### RNA-Seq identifies Fyn driven activation of Stat3, metabolic, oxidative stress, and inflammatory signaling pathways

To identify altered pathways and potential downstream effectors of Fyn signaling in neurodegeneration we performed bulk RNA-Seq on RNA extracted from 3 dpf and 5 dpf control *elavl3:Gal4* and *elavl3:Gal4; UAS:fynY531F* larva (Figure 5; n=4 for each condition and genotype). The top 50 genes with highest changes in gene expression in 5 dpf vs. 3 dpf *elavl3:Gal4; UAS:fynY531F* larva included serine proteases *prss1* and *prss59.1*, and cathepsins *ctsbb* and *ctsl*, which are involved in proteolysis and metabolism in cytokine-related neuroinflammation (Ha et al. 2022; Jiang et al. 2022). Top genes in 5 dpf *elavl3:Gal4; UAS:fynY531F* vs. control *elavl3:Gal4* included the ortholog of activated macrophage gene *acod1/irg1* immune responsive gene *irg1l* and the Stat3 target *timp4.1* (Fig. 5A). KEGG pathway analysis (Fig. 5B) indicated changes in electrical transmission, protein translation, and metabolism of carbon, tryptophan, and fatty acids, and the PPAR peroxisome proliferator-activated receptor pathway, which mediates inactivation of NF-κB during the inflammatory response (Korbecki, Bobinski, and Dutka 2019). Volcano plots (Fig. 5C) comparing 3 dpf vs 5dpf *elavl3:Gal4; UAS:fynY531F* and 5 dpf *elavl3:Gal4; UA*S:fynY531F vs control show reduced expression of the neuroprotective genes *apoea*, *sod3a*, and *oxsr1a*. Apolipoprotein E (*apoea*) is essential for maintaining cholesterol homeostasis and neuronal function and is a primary risk factor and therapeutic target in AD (Williams, Borchelt, and Chakrabarty 2020). Superoxide dismutase 3 (*sod3a*) encodes a key antioxidant enzyme protecting cells from oxidative stress-induced damage, a hallmark of neurodegeneration (Olufunmilayo, Gerke-Duncan, and Holsinger 2023).

**Figure 5.**
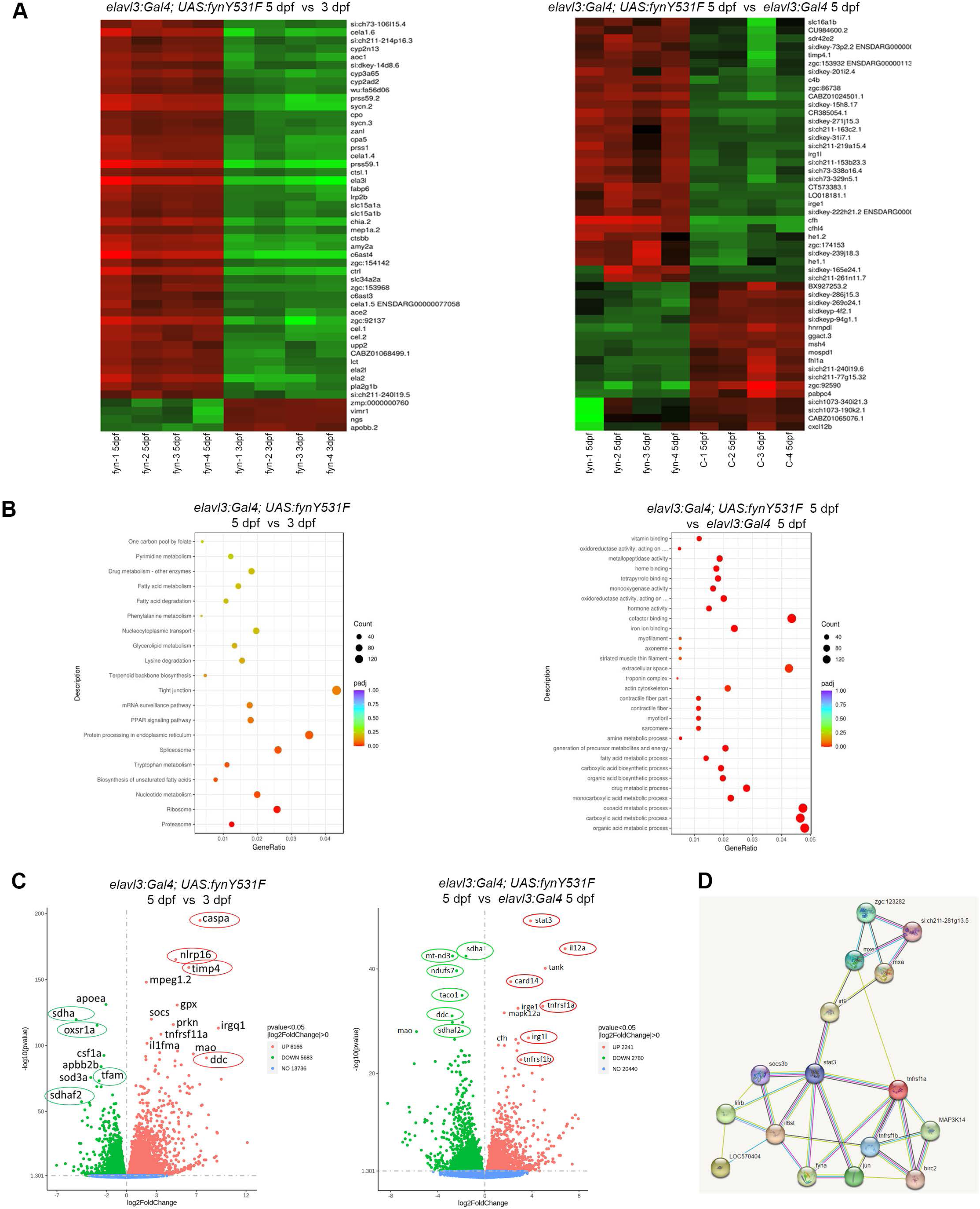
Transcriptome analysis reveals FynY531F signaling alters neuroprotective, oxidative and metabolic pathways and identifies Stat3 as a downstream Fyn effector. **A** Heatmaps of highest gene expression changes comparing bulk RNA Seq from 5 dpf vs 3 dpf *elavl3:Gal4; UAS:FynY531F* larva (n=4; left heat map), and 5 dpf *elavl3:Gal4; UAS:FynY531F* vs 5 dpf *elavl3:Gal4* (n=4; right heat map). **B** Pathway analysis reveals altered metabolic pathways in 5 dpf vs 3 dpf *elavl3:Gal4; UAS:FynY531F* (left plot) and 5 dpf *elavl3:Gal4; UAS:FynY531F* vs 5 dpf *elavl3:Gal4* (right plot). **C** Volcano plots showing up and downregulated genes in 5 dpf vs 3 dpf *elavl3:Gal4; UAS:FynY531F* (left plot) and 5 dpf *elavl3:Gal4; UAS:FynY531F* vs 5 dpf *elavl3:Gal4* (right plot). Nuclear encoded mitochondrial genes *sdha*, *sdhaf2* and *taco1* are reduced. Elevated genes include *stat3*, *caspa*, *tngrsf1a, b* and *irg1l*. **D** String network analysis identifies Fyn interactions with Tnf-α and Stat3 signaling pathways in *elavl3:Gal4; UAS:FynY531F* larval transcriptome.

Oxidative stress responsive 1 homolog A (*oxsr1a*) is essential for cellular resistance to oxidative stress and its downregulation has been show to precede the onset of neurodegeneration (Volkert and Crowley 2020).

Consistent with oxidative stress as a factor in driving neurodegeneration, there was also a reduction in genes required for mitochondrial function (*taco1, mt-nd3*, *ndufs7*, *mao, tfam*) (Oktay et al. 2020; Richman et al. 2016) and oxidative phosphorylation (*sdha, sdhaf2)* (Fullerton et al. 2020). Elevated metabolic pathways identified by KEGG analysis above suggest a compensatory mechanism for an energy deficit resulting from reduced mitochondrial function. Decreased Dopa decarboxylase gene *ddc* suggest a disruption in dopaminergic neurotransmitter metabolism (Ramesh and Arachchige 2023), that correlates with loss of dopaminergic neurons in *elavl3:Gal4; UAS:fynY531F* larvae. Significant elevation of genes related to apoptosis or programmed cell death were not detected, although there was a reduction in caspase3 and apoptosis-related cysteine peptidase in 5 dpf vs 3dpf *elavl3:Gal4; UAS:fynY531F* larvae. Together these results suggest Fyn signaling may contribute to neurodegeneration through disruption of neuroprotective mechanisms and energy production.

Upregulated pathways consistent with cytokine RT-qPCR gene expression analyses described above included elevated expression of inflammatory signaling components (*nlrb16, caspa*, *card14*), and microglia activation (*irg1l*) (Fig. 5C). A significant increase was also detected in components of the Signal transducer and transcription activator pathway, including *stat3*, *timp2b*, *timp4.1*, *socs3* (Fig. 5C) and Tnf-α receptors *tnfrsf1a*, *b* which have been shown to be a direct transcriptional target of Stat3 in breast cancer cells (Egusquiaguirre et al. 2018). To reveal potential protein-protein interactions, String network analysis was performed and identified connections between Fyn and Stat3 signaling (Fig. 5D), providing additional evidence for Stat3 as a potential novel downstream effector of Fyn. Together these gene expression analyses indicate Fyn driven neurodegeneration may be due to defective neuroprotective mechanisms that correlate with oxidative stress, inflammatory cytokine production, and Stat3 pathway activation.

### Stat3 is a novel Fyn downstream effector driving dopaminergic neuron loss and cytokine induction

Differential gene expression in the FynY531F *elavl3:Gal4; UAS:fynY531F* model identified Stat3 as a potential downstream pathway activated by Fyn signaling. In cultured human melanoma cells Fyn kinase has been shown to phosphorylate Stat3 on Tyr705 (Tang et al. 2020). To test whether Fyn signaling drove Stat3 activation in the zebrafish Fyny531F model, western blotting of wild type +/+, control *elavl3:Gal4*, and *elavl3:Gal4; UAS:fynY531F* larvae was performed with an anti-Stat-Y705-PO antibody (Fig. 6A). At 3 dpf and 5 dpf elevated levels of Stat-Y705-PO were detected in *elavl3:Gal4; UAS:fynY531F* larvae compared to wildtype and control samples (Fig. 6A, B). Treatment of *elavl3:Gal4; UAS:fynY531F* from 3 dpf to 5 dpf with 10μM Src family inhibitor Saracatinib inhibited Stat-Y705-PO levels, which remained similar to the level detected in control *elavl3:Gal4* larvae (Fig. 6A, B). These results show Fyny31F signaling drives Stat3-Y705 phosphorylation that is dependent on Fyn kinase activity.

**Figure 6.**
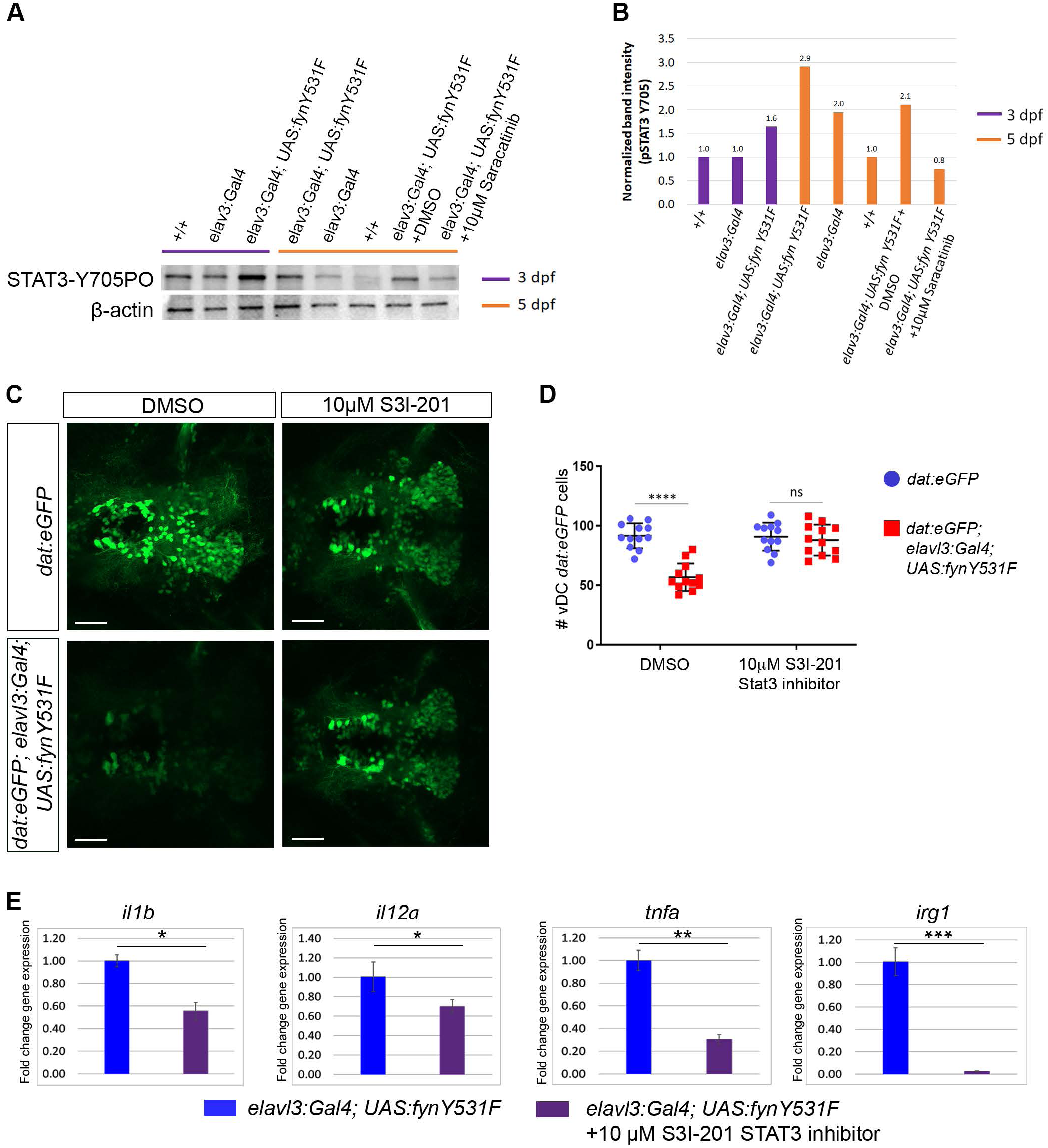
Stat3 inhibition suppresses FynY531F driven dopaminergic neuron degeneration and induction of *il1b*, *il12a*, *tnfa* and *irg1* expression. **A** Western blot of 3 dfp and 5 dpf +/+, control *elavl3:Gal4* and *elavl3:Gal4; UAS:FynY531F* larval extracts probed with anti-Stat3-Y705-PO antibody shows increased Stat3-Y705 phosphorylation in *elavl3:Gal4; UAS:FynY531F* larvae. *elavl3:Gal4; UAS:FynY531F* larvae treated from 3 dpf to 5 dpf with 10 μM Src family inhibiot Saracatinib show reduced Stat3-Y705 phosphorylation. Blot was probed with anti-GAPDH and anti-βactin as protein loading controls. **B** Quantification of band intensities on Stat3-Y750-PO Western blot. **C** Live confocal imaging of vDC eGFP dopaminergic neurons in control *dat:eGFP* and *dat:eGFP;elav:Gal4; UAS:fynY531F* 5 dpf larvae show rescue of dopaminergic neuron loss after treatment with 10 μM Stat3 inhibitor S3I-201. **D** Quantification of vDC eGFP dopaminergic neuron number in control *dat:eGFP* and *dat:eGFP;elav:Gal4; UAS:fynY531F* 5 dpf larvae after treatment with 10 μM Stat3 inhibitor S3I-201 (n=12). Analysis was performed using two way ANOVA with Tukey’s multiple comparison. **E** RT-qPCR of *il1b*, *il12a*, *tnfa*, and *irg1* on RNA extracts from untreated and 10μM Stat3 inhibitor S3I-201 treated *elavl3:Gal4; UAS:fynY531F* 5 dpf larvae (n= 3 biological replicates for each genotype). RT-qPCR analysis was performed with two-tailed unpaired Student’s t-test. Bars represent mean +/- SEM. ns, not significant; * p< 0.05; ** p<0.01; *** p< 0.001. Scale bars 100μm.

To determine whether dopaminergic neuron loss in the FynY531F model was dependent on Stat3 activation, *elavl3:Gal4; UAS:fynY531F* larvae were treated with 10μM of the Stat3 inhibitor S3I-201 followed by quantification of the vDC dat:eGFP cluster in the ventral diencephalon (Fig. 6C, D). Stat3 inhibitor treatment from 3 dpf to 5 dpf suppressed *dat:eGFP* neuron loss in *elavl3:Gal4; UAS:fynY531F* larvae (Fig. 6C). The number of *dat:eGFP* positive cells in treated *elavl3:Gal4; UAS:fynY531F* larvae was not significantly different from untreated control (p=0.99) or treated control larvae (p=0.92) (Fig. 6D). Together with the results shown above, this analysis indicates Stat3 is a downstream effector of Fyn signaling that mediates dopaminergic neuron loss.

We next examined whether elevated cytokine levels and microglia activation in *elavl3:Gal4; UAS:fynY531F* larvae were dependent on Stat3 activation. RT-qPCR on *elavl3:Gal4; UAS:fynY531F* larvae treated with 10μM Stat3 inhibitor S3I-201 showed treatment with the Stat3 inhibitor suppressed elevated levels of cytokines *il1b* (p<0.05), *il12a* (p<0.04), and *tnfa* (p<0.001), and microglia activation marker *irg1* (p>0.0001). These results are consistent with Fyn signaling driving Stat3 activation and induction of inflammatory cytokine expression that correlates with dopaminergic neuron loss and microglia actvation.

### FynY531F induced dopaminergic neuron loss and cytokine elevation depend on NF-κB inflammatory signaling

The results presented above indicate Fyn signaling drives induction of inflammatory cytokines *il1b* and *il12a*, that are known to be expressed by activated microglia via the NF-κB pathway (Wang et al. 2015). To test whether FynY531F driven dopaminergic neuron loss was dependent on the NF-κB inflammatory signaling pathway, *dat:eGFP*; *elavl3:Gal4; UAS:fynY531F* larvae were treated at 3 dpf with NF-κB and Caspase 1 inhibitors, and collected at 5 dpf for live imaging and quantification of vDC dopaminergic neurons. Treatment of *dat:eGFP*; *elavl3:Gal4; UAS:fynY531F* larvae with 0.5 μM NF-κB inhibitor CAPE suppressed vDC dopaminergic neuron loss (Fig. 7A). The numbers of eGFP positive vDC neurons in treated larvae showed no significant difference compared to control *dat:eGFP* larvae (p=0.96) (Fig. 7B). Treatment of *dat:eGFP*; *elavl3:Gal4; UAS:fynY531F* larvae with 20 μM Caspase-1 inhibitor Ac-YVAD-cmk also suppressed vDC dopaminergic neuron loss compared to control larvae (Fig. 7 A,B p=0.08). These results determine that constitutive FynY531F kinase signaling driving dopaminergic neuron loss is mediated through activation of the inflammatory signaling pathway NF-κB/Caspase-1.

**Figure 7.**
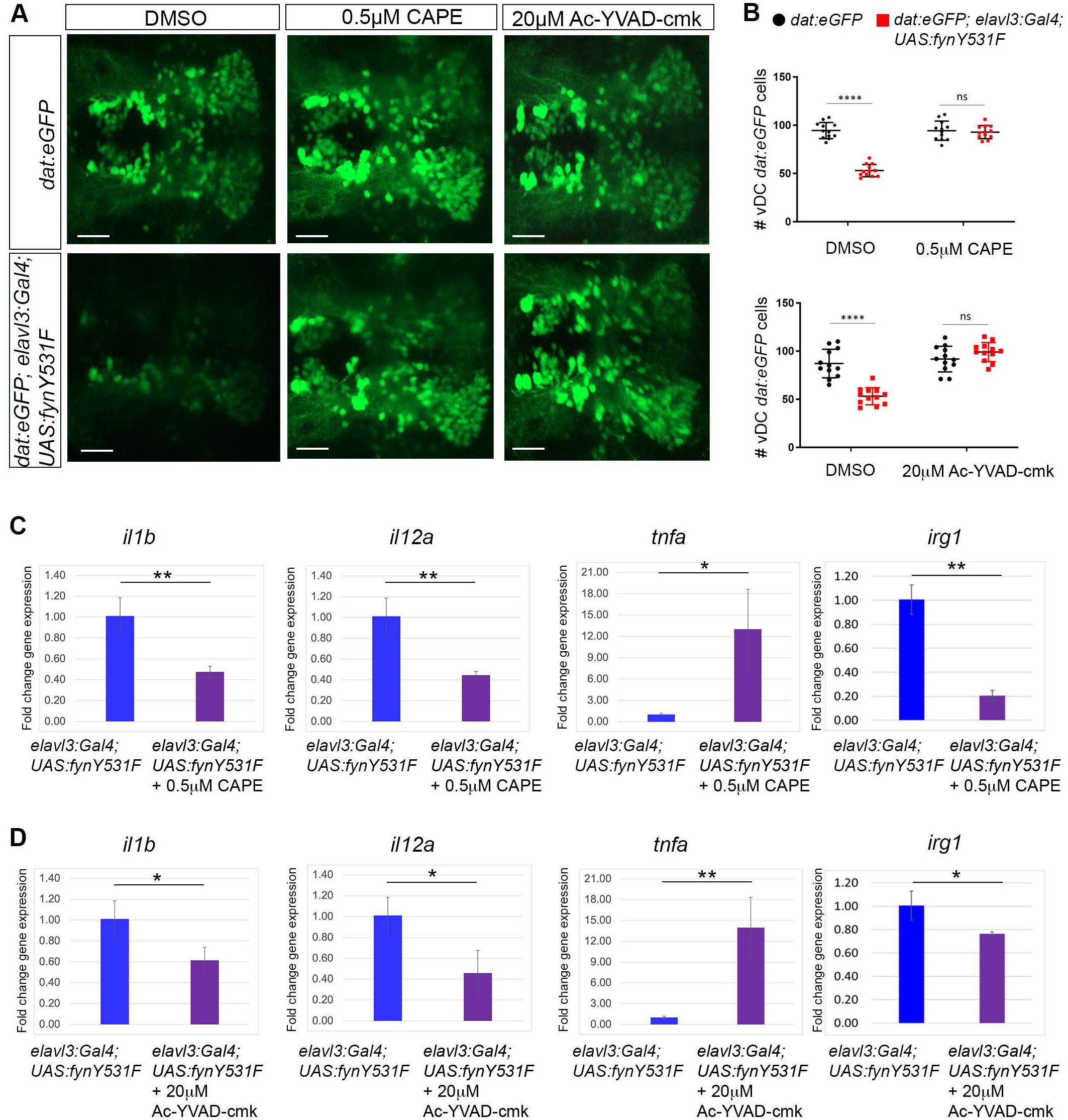
NF-κB and Caspase 1 inhibition suppresses FynY531F driven dopaminergic neuron loss and induction of *il1b*, *il12a* and *irg1* expression. **A** Live confocal imaging of vDC eGFP positive dopaminergic neurons in control *dat:eGFP* and *dat:eGFP;elav:Gal4; UAS:fynY531F* 5 dpf larvae show rescue of dopaminergic neuron loss after treatment with 0.5µM Caffeic acid phenethyl ester (CAPE-NF-κB inhibitor) and 20µM Ac-YVAD-cmk (Caspase 1 inhibitor). **B** Quantification of vDC eGFP+ cells (n=12). Analysis was performed using two way ANOVA with Tukey’s multiple comparison. **C, D** RT-qPCR showing significantly reduced levels of *il1b*, *il12a* and *irg1* after treatment of *dat:eGFP;elav:Gal4; UAS:fynY531F* larvae with 0.5µM CAPE and 20µM Ac-YVAD-cmk inhibitors. *tnfa* was increased after treatment with 0.5µM CAPE and 20µM Ac-YVAD-cmk inhibitors. (n= 3 biological replicates for each genotype and condition). RT-qPCR was analysis was performed with Two-tailed unpaired Student’s t-test. Bars represent mean +/- SEM. ns, not significant; * p<0.05; ** p<0.01. Scale bars 100μm.

RT-qPCR was used to determine whether FynY531F *elavl3:Gal4; UAS:fynY531F* elevated inflammatory cytokine expression was dependent on the NF-κB/Caspase-1 pathway. Treatment of *elavl3:Gal4; UAS:fynY531F* larvae with NFkΒ inhibitor 0.5 μM CAPE suppressed the increase in *il1b*, *il12a*, and *irg1* expression (Fig. 7C). The result with Caspase-1 inhibitor 20 μM Ac-YVAD-cmk was similar, with a slightly less significant difference in expression level compared to untreated *elavl3:Gal4; UAS:fynY531F* larvae (Fig. 7D). Neither inhibitor suppressed elevated *tnfa* expression in *elavl3:Gal4; UAS:fynY531F* larvae. Together, these results suggest microglia and NF-κB/Caspase-1 pathway activation occur in response to an external inflammatory signal, which may originate in degenerating dopaminergic neurons.

### Stat3 and NFκΒ synergize in Fyn driven dopaminergic neuron loss

To examine the interaction of Stat3 and NF-κB pathways in Fyn driven dopaminergic neuron loss *elavl3:Gal4; UAS:fynY531F* larvae were treated either alone or with a combination of Stat3 and NF-κB inhibitors (Fig. 8A). Dual treatment of larva with 10 μM S31-301 Stat3 inhibitor and 0.5 μM CAPE NF-κB inhibitor resulted in toxicity. Therefore, the amount of each inhibitor was reduced by one half to test for interaction of the two pathways. Dual treatment of larva with 5 μM S31-301 and 0.25 μM CAPE suppressed the loss of *dat:eGFP* signal to a significantly higher level than either inhibitor alone (Fig. 8A, B) (10 μM S31-301 vs. 5 μM S31-301 plus 0.25 μM CAPE, p=0.003; 0.5 μM CAPE vs. 5 μM S31-301 plus 0.25 μM CAPE, p=0.002). These results indicate Fyn activates Stat3 and NF-κB pathways that act synergistically to drive dopaminergic neurodegeneration.

**Figure 8.**
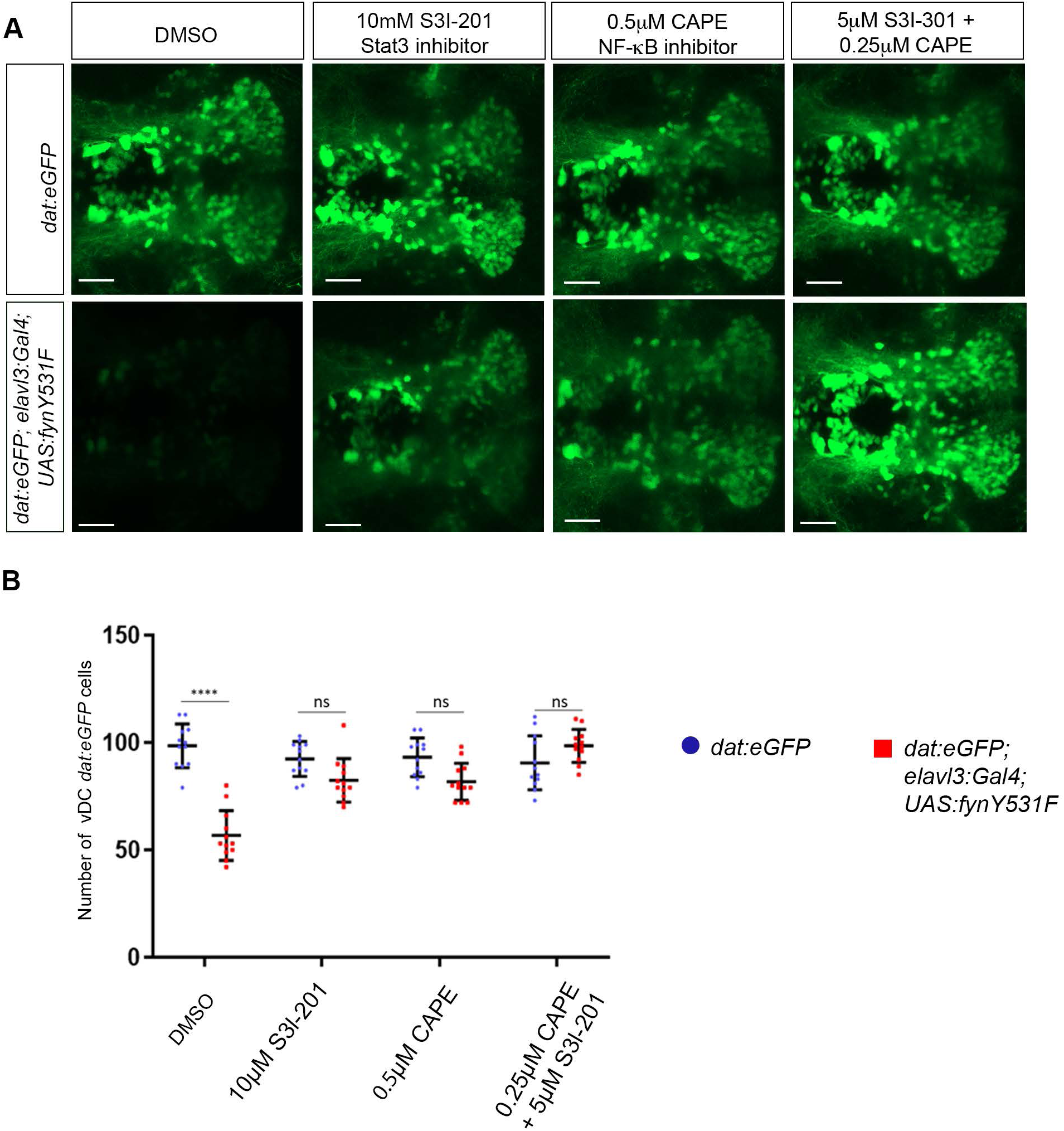
Dual inhibition of NF-κB and Stat3 acts synergistically to suppresses FynY531F driven dopaminergic neuron degeneration. **A** Live confocal imaging of vDC eGFP dopaminergic neurons in control *dat:eGFP* and *dat:eGFP;elav:Gal4; UAS:fynY531F* 5 dpf larvae after treatment with mock DMSO, 10 μM Stat3 inhibitor S3I-201, 0.5 NF-κB inhibitor CAPE, and 5 μM Stat3 inhibitor S3I-201 plus 0.25 NF-κB inhibitor CAPE. **B** Quantification of vDC eGFP dopaminergic neuron number in control *dat:eGFP* and *dat:eGFP;elav:Gal4; UAS:fynY531F* 5 dpf larvae after treatment mock DMSO, 10 μM Stat3 inhibitor S3I-201, 0.5 NF-κB inhibitor CAPE, and 5 μM Stat3 inhibitor S3I-201 plus 0.25 NF-κB inhibitor CAPE (n=12). Analysis was performed using two way ANOVA with Tukey’s multiple comparison. Bars represent mean +/- SEM. **** p<0.0001. Scale bars 100μm.

## Discussion

Here we describe a novel zebrafish model of activated Fyn signaling which demonstrates a neural specific role for Fyn driven neurodegeneration. The zebrafish Fyn model provides *in vivo* evidence consistent with a role for Fyn signaling in the pathology of neurodegenerative disorders and reports of elevated Fyn signaling in patient brain tissue (Low et al. 2021; Panicker et al. 2019; Guglietti et al. 2024). In the zebrafish Fyn model activated Fyn drives inflammatory cytokine production, leading to dopaminergic neuron loss, mitochondrial dysfunction and microglia activation. Chemical inhibition demonstrates Fyn drives neurodegeneration through activation of Stat3 and NF-κB/Caspase 1, which synergize in dopaminergic neuron loss. While both Stat3 and NF-κB were required for induction of *il1b* and *il12a*, *tnfa* elevation was only dependent on Stat3 activation. Our study suggests a model in which Fyn drives production of *tnfa* through activation of Stat3 signaling in neurons. Release of *tnfa* may contribute to microglia activation, driving production of inflammatory cytokines *il1b* and *il12a*, which in turn stimulate a sustained neuroinflammatory response correlated with mitochondrial dysfunction and neuron loss (Graphical Abstract).

Activated Fyn signaling led to dopaminergic neuron loss over the course of 3 days to 5 days post fertilization, after the onset of neural differentiation in the larval brain. Gene expression changes coincident with this time frame indicate disruption of neuroprotective mechanisms and mitochondrial energy production. An increase in apoptosis related genes was not detected, indicating dopaminergic neuron loss was not linked to programmed cell death pathways. These observations are distinct from reports of Src family/Fyn kinase phosphorylation of PKC-δ leading to rat dopaminergic neuron oxidative stress and cell death through caspase-mediated apoptosis (Kaul et al. 2005; Saminathan et al. 2011). The increase in mitochondrial signal in dopaminergic cell bodies in 3 dpf larva, before loss of dat:eGFP labeled dopaminergic neurons, suggests mitochondrial dysfunction may be a driving factor in neuron loss. Mitochondrial dysfunction is a well-established factor contributing to neurodegeneration (Klemmensen et al. 2024). In our model, Fyn signaling resulted in accumulation of dat:mitoRFP in the cell bodies of dopaminergic neurons and downregulation of genes crucial for mitochondrial respiratory complex function (*sdha, sdhaf2, mt-nd3, ndufs7, taco1*) (Borna et al. 2024; Machado et al. 2016; Richman et al. 2016; Oktay et al. 2020; Fullerton et al. 2020) and biogenesis (*tfam*) (Kang, Kim, and Hamasaki 2007; Stiles et al. 2016). In addition, mitochondrial dysfunction may be a direct result of Fyn kinase targeting mitochondrial proteins for phosphorylation. Mitochondrial elongation factor EF-Tu_mt_ is phosphorylated on Tyr266 by Fyn and c-Src kinases, which was shown to inhibit translation *in vitro* (Koc, Hunter, and Koc 2023). Elevated Fyn signaling may disrupt oxidative energy metabolism through negative regulation of mitochondrial protein synthesis and homeostasis. Alternatively, mitochondrial aggregation in dopaminergic cell bodies may result from a defect in overall reduced organelle transport in axons.

Transcriptome analysis of the zebrafish Fyn model identified Stat3 as a novel downstream effector of Fyn signaling in neurodegeneration. Like Fyn, Stat3 signaling has been implicated in neurodegeneration. Fyn and Stat3 were identified as potential AD biomarkers in young APOE ε4 individuals (Roberts et al. 2021). *in vitro* analyses demonstrated Stat3 and Fyn inhibitors suppressed AD related phenotypes of LPS-induced neuroinflammation, tau phosphorylation, and Aβ secretion (Roberts et al. 2021). Stat3-PO is observed in AD patient postmortem brain tissue, in mouse APP/PS1 AD model brain, is mediates in vitro Ab induced neuron cell death (Wan et al. 2010). Astrogliosis in the mouse APP/PS1 AD model is dependent on Stat3 (Reichenbach et al. 2019; Ben Haim et al. 2015). Stat3-Y705-phosphorylation has been shown to regulated by Src kinases in cultured tumor cells (Silva 2004; Ram and Iyengar 2001; Garcia et al. 2001), and Fyn is required for Stat3-PO activation in T cell receptor signaling and T cell differentiation (Qin et al. 2024). Our model suggests *in vivo*, Fyn and Stat3 function in the same pathway to drive neuroinflammation and neurodegeneration, with Stat3 activation dependent on Fyn signaling.

Dual chemical inhibition revealed Stat3 and NF-κB pathways synergize in dopaminergic neuron loss in the Fyn model. Chemical inhibition of Stat3 significantly suppressed all three elevated cytokines, *il1b*, *il12a*, and *tnfa,* supporting a role for Stat3 in induction of multiple inflammatory signaling pathways and microglia activation. While inhibiting NF-κB and Caspase-1 suppressed expression of *il1b* and *il12a*, it did not suppress *tnfa* induction, suggesting a temporal and spatial separation of activation of the Stat3 and NF-κB pathways. However, Stat3 gene targets elevated in the Fyn neurodegeneration model included *tnfrsf1a*, which encodes a receptor for TNFα and is associated with activation of the NF-κB pathway in breast cancer (Egusquiaguirre et al. 2018). Stat3 and NF-κB pathways have also previously been shown to interact in hepatic cells. Stat3 and RelA/p65 formed a complex in HepG2 hepatoblastoma cells stimulated with IL-1 and IL-6 (Hagihara et al. 2005), and NF-κB RelA/p65 homodimers were shown to cooperate with Stat3 in Hep3B cells in response to IL-1 (Yoshida et al. 2004). In cultured microglia, Fyn activation was shown to lead to cytokine production through activation of PKC-δ/NF-κB/Caspase 1 inflammasome signaling (Panicker et al. 2015; Panicker et al. 2019). In our *in vivo* model, the possibility of Fyn directly activating the NF-κB pathway in neurons cannot be excluded, nor can indirect activation of Stat3 in microglia leading to stimulation of the NF-κB pathway. Microglia are known to produce TNFα when activated through the JAK/Stat pathway (Yin et al., 2018). Distinguishing these possibilities would require cell type specific inhibition to determine whether Stat3 and NF-κB/Caspase 1 pathways are functioning in dopaminergic neurons, microglia or both. Nevertheless, our findings suggest an interplay between Fyn-driven cytokine production in neurons and activated microglia, potentially creating a feedback loop that fuels persistent neuroinflammation and neurodegeneration.

Our *in vivo* zebrafish model of neural Fyn signaling reveals Fyn drives dopaminergic loss through inflammatory cytokine production and microglia activation, and unveils Stat3 as a potential novel downstream Fyn effector. The synergistic effect of Stat3 and NF-kB inhibition in suppressing Fyn driven dopaminergic loss support the contribution of both pathways in mediating Fyn neurodegeneration. Further investigation is necessary to identify the role of Stat3 and its cellular targets in driving dopaminergic neuron loss and mitochondrial dysfunction in response to Fyn signaling.

## Materials and Methods

### Ethics Declarations and approval for animal experiments

Use of zebrafish for research in this study was performed according to the Guidelines for Ethical Conduct in the Care and Use of Animals (APA 1986), and carried out in accordance with Iowa State University Animal Care and Use Committee IACUC-21-281 and IBC-21-117 approved protocols. All methods involving zebrafish were in compliance with the American Veterinary Medical Association (2020), ARRIVE (Percie du Sert et al. 2020) and NIH guidelines for the humane use of animals in research.

### Zebrafish maintenance and transgenic strains

Zebrafish *Danio rerio* used in this study were housed in the Roy J. Carver Charitable Trust Zebrafish Facility at Iowa State University. Fish were maintained at 28.5°C on an Aquaneering aquaculture system with a 14-hour light/10-hour dark light/dark cycle. Embryos were collected from natural spawning and were raised in E3 embryo media (13 mM NaCl, 0.5 mM KCL, 0.02 mM Na2HPO4, 0.04 mM KH2PO4, 1.3 mM CaCl2, 1.0 mM MgSO4, 4.2 mM NaHCO3, pH 7.0). Wild type WIK zebrafish were obtained from the Zebrafish International Resource Center (https://zebrafish.org/home/guide.php). Transgenic zebrafish previously described used in this study include dopaminergic neuron eGFP and mito-RFP reporters *Tg*(*dat:eGFP) (dat:eGFP)* (Xi et al. 2011) and *Tg(dat:tom20 MLS-mCherry) (dat:mitoRFP)* (Noble et al. 2015), and the pan-neuronal Gal4 driver *Tg(elavl3:Gal4-VP16)nns6* (Kimura, Satou, and Higashijima 2008).

### Isolation of *Tg(UAS:fynY531F)* transgenic lines

The 1614 bp *fyna* (ENSDARG00000011370) wild type and Y531F mutant cDNAs were amplified by RT-PCR from RNA isolated from 3 dpf embryos using primers listed in Table 1. The forward and reverse primers incorporated KpnI and BssHII restriction enzyme sites, respectively. 500ng RNA was used as a template for reverse transcription with Superscript II (Invitrogen 11752) followed by amplification with KOD polymerase (Sigma-Aldrich 71842). Mutant *fynY531F* was directionally cloned into the transposon vector *Tol2(14XUAS, gcry1:eGFP)* (Balciuniene et al. 2013) to build the *Tol2 (UAS:fynY531F; gcry1:eGFP)* construct. *Tol2* transposase mRNA was *in vitro* transcribed from 1 ug linear *pT3TS-Tol2* plasmid (Balciunas et al. 2006) to generate capped, polyadenylated mRNA using T3 mMessage mMachine High Yield Capped RNA transcription kit (ThermoFisher AM 1348). *Tol2* mRNA was purified using the ZYMO RNA Clean & Concentrator-5 kit (ZYMO R1015) and eluted in RNase-free water. *Tg(Tol2<UAS:fynY531F>)is89 and Tg(Tol2<UAS:fynY531F>)is90* lines were isolated by coinjection of 50 pg *Tol2* mRNA and 25 pg transposon vector into 1-cell WIK embryos. 5 *Tg(Tol2<UAS:fynY531F>)* F0 adults were screened to identify two independent founders transmitting a *Tol2* integration through the germline. Individual F1 adults were used to generate F2 fish, and a single F2 adult showing Mendelian segregation of transmitted alleles was outcrossed to WIK to establish the independent lines *Tg(Tol2(14XUAS:fynaY531F, gcry1:eGFP>)is89* and *Tg(Tol2(14XUAS:fynaY531F, gcry1:eGFP>)is90*.

**Table 1.**
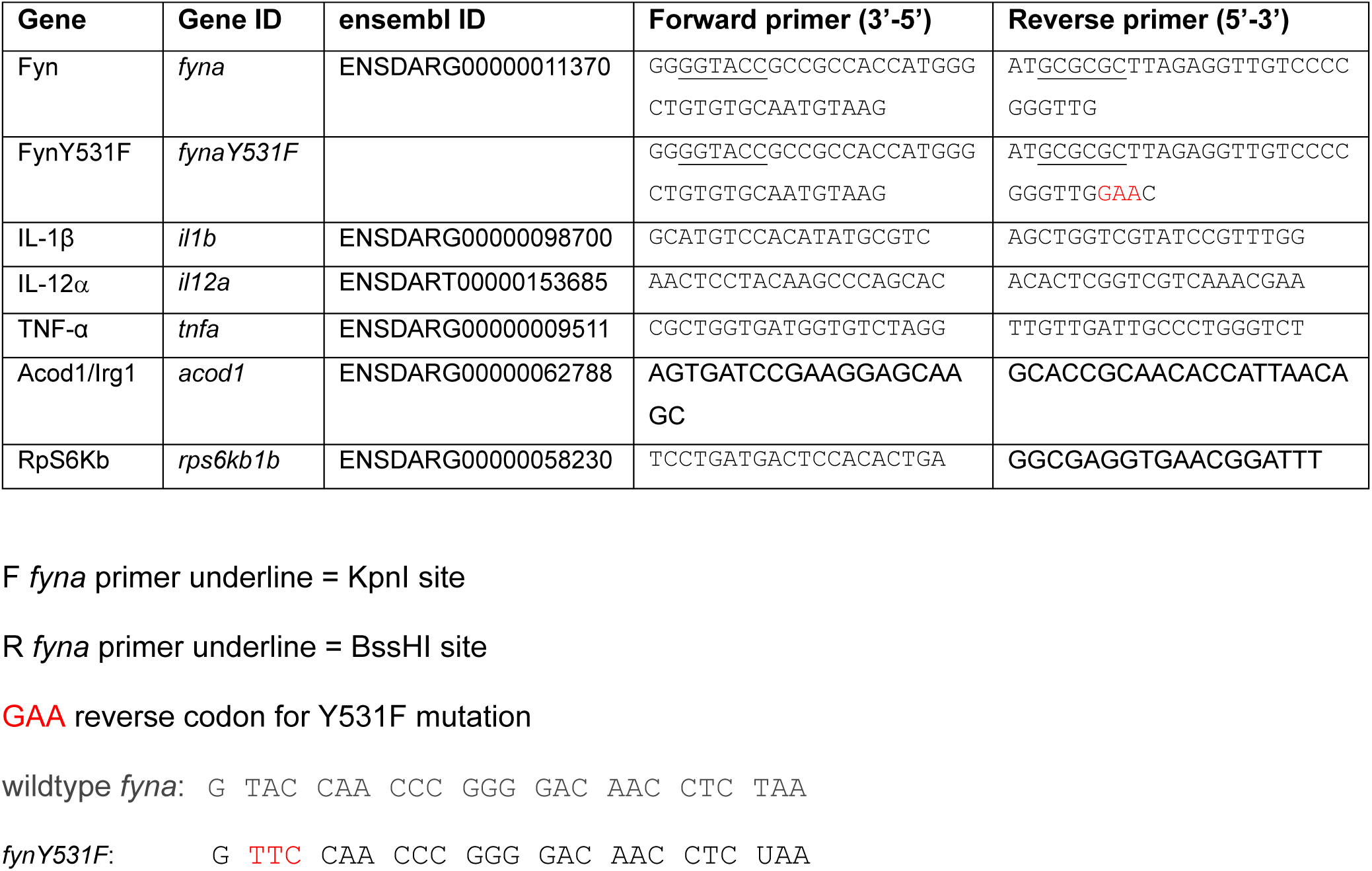
Oligonucleotide primers used in this study.

### Behavioral Analysis

Zebrafish larvae were monitored for swimming behavior using a ZebraBox monitoring system and ZebraLab software (ViewPoint Behavior Technology). 4 dpf larvae were placed in a 48-well plate in E3 embryo media. At 5dpf larvae were placed in the viewing chamber and acclimated for 30 minutes before recording. Locomotion was recorded for 5 hours under alternating light/dark conditions in 15 minute intervals. Larval movement, velocity, and distance were recorded each minute and the data analyzed and plotted with ZebraLab software.

### Chemical inhibitor treatment assays

For chemical inhibitor assays, 10 larvae of each genotype at 3 day post-fertilization (dpf) were placed in a well in a 6-well plate in 3 mL embryo media containing 0.003% 0.2 M phenylthiourea (PTU) and designated inhibitor diluted to the final concentration, or DMSO as a vehicle control. The larvae were placed in a 28.5°C incubator. The chemical inhibitor or DMSO solution was replaced with fresh solution in the morning of 4 dpf and 5 dpf. Caspase-1 inhibitor - InvitroFit™ Ac-YVAD-cmk (Garcia-Calvo et al. 1998) (InvivoGen inh-yvad) was dissolved in DMSO and used at 20µM final working concentration. NF-κB inhibitor Caffeic acid phenethyl ester CAPE (Natarajan et al. 1996) (Apexbio B1644) in DMSO was used at 0.5µM final working concentration. Src Family Kinase inhibitor Saracatinib AZD0530 (Green et al. 2009) (MedChemExpress HY-1-234) in DMSO was used at 20µM final working concentration. Stat3-PO inhibitor S31-301 was used at a final concentration of 10 µM. For dual inhibitor assays, CAPE and S31-301 were used a final concentration of 0.25µM and 5 µM, respectively.

### RNA isolation, RT-quantitative PCR, RNA-Seq

Reverse Transcription-quantitative PCR experiments were designed and carried out according to updated MIQE guidelines (Bustin et al. 2009; Taylor et al. 2019). For RT-qPCR, 30 5 dpf larvae per biological replicate were placed in Zymo Research DNA/RNA Shield (R1100-50) and homogenized using a disposable pestle. RNA extraction was performed according to the manufacturer’s instructions. 500 ng total RNA was used as template with the Luna Universal One-step RT-qPCR kit (New England Biolabs, E3005L). For each condition 3 biological replicates with two technical replicate RT-qPCR reactions were run on a CFX96 Connect Real-Time System (BioRad). The number of *il1b*, *il12a*, *tnfa*, and *irg1* transcripts was quantified using the Comparative CT method (ΔΔCT) and normalized using *rps6kb1b* reference gene. The oligonucleotide primers used for RT-qPCR are listed in Table 1. For RNA-Seq total RNA was extracted as above. Library construction, PE150 next generation sequencing, and differential gene expression was performed at Novogene using DeSeq2. Differential gene expression data were analyzed with online software for heat map generation and KEGG analysis (http://bioinformatics.sdstate.edu/idep96/) and STRING analysis (https://string-db.org/).

### Western Blotting

Protein extracts were generated from zebrafish larvae placed in lysis buffer 50mM Tris-HCl pH 7.5, 150mM NaCl, 5mM MgCl_2_, 1% Triton X-100, 0.5% NP-40 containing 1X Halt Protease Inhibitor Single-use cocktail (Thermo Scientific 78430) and ground with a disposable pestle. SDS-PAGE of protein extracts was performed with a BioRad Mini-Protean gel system using Mini-PROTEAN TGX Precast 4-15% polyacrylamide gels (BioRad 4561083) and blotted to Biorad Immun-Blot LF PVDF membrane (BioRad 1620174). Blots were blocked with Blot-Qualified BSA (Sigma A7906) and probed with the following primary antibodies: anti-416-PO Srk Kinase Family rabbit polyclonal (Cell Signaling Technology 6943T) at 1:1000, anti-Stat3-Y705PO XP rabbit monoclonal antibody (Cell Signaling Technology 9145), and anti-GAPDH mouse monoclonal antibody (Proteintech 60004-1-Ig). Goat anti-Mouse IgG Horse Radish Peroxidase (Invitrogen 31430) and Goat anti-Rabbit IgG Horse Radish Peroxidase (Invitrogen 31460) secondaries were used at 1:10,000 dilution. Western blots were developed with SuperSignal West Dura Extended Duration Substrate (Thermo Scientific 34075) and imaged on an iBright system (ThermoFisher FL1500).

### Zebrafish immunolocalization and live confocal imaging

Zebrafish larvae fixation, embedding, sectioning and immunolabeling was as described previously (Schultz et al. 2018). Embryos were incubated in 0.003% 1-phenyl 2-thiourea (Sigma P7629) in E3 embryo media to inhibit pigment synthesis. Larvae were euthanized in 160ug/ml Ethyl 3-aminobenzoate methanesulfonate and fixed in 4% paraformaldehyde (PFA) overnight at 4°C or in 2% trichloroacetic acid for 3 hours at room temperature. Whole mount fixed larvae were labeled to visualize macrophages and microglia with the mouse hybridoma supernatant 4C4 (ECACC 92092321, A. Dowding, King’s College London, UK) a gift from Dr. Diana Mitchell, University of Idaho, at a 1:100 dilution. Rabbit polyclonal anti-acetylated alpha tubulin antibody (Invitrogen PA5-105102) was used at 1:500. Secondary antibodies Alexa Flour 594 goat anti-mouse IgG (H+L) (Thermo Fisher A11005) and Alexa Flour 488 goat anti-rabbit IgG (H+L) (Thermo Fisher A11008) were used at 1:500. For live confocal imaging of *dat:eGFP* and *dat:mito-RFP* dopaminergic neurons larvae were mounted in 1.2% low-melting point agarose in E3 embryo media containing 160ug/ml Ethyl 3-aminobenzoate methanesulfonate (Tricaine MS-222; Sigma-Aldrich 886-86-2) Tricaine anesthetic. Larva and immunolabeled tissues were imaged on a Zeiss LSM 800 laser scanning confocal microscope. Diencephalic dopaminergic vDC clusters images were acquired by maximum projection of Z-stacks taken at 2 μm sections.

### Quantification and statistical analyses

*dat:eGFP* positive cell counts in the larval diencephalic dopaminergic neuron dVC cluster were quantified from 12 larvae for each genotype and condition. Quantification was analyzed with Two-way ANOVA followed by Tukey’s multiple comparison test. RT-qPCR to determine gene expression levels was performed on three independent pools of 30 larvae for each condition. 4C4 microglia counts, mitoRFP/GFP cell counts, and RT-qPCR data were analyzed using two-tailed unpaired Student’s t-test with mean ± s.e.m. Gene expression data was analyzed using DeSeq2 (Novogene). Statistical significance was considered to be present when the p-value was less than 0.05 and is indicated as *p < 0.05, **p < 0.01, ***p < 0.001 or ****p < 0.0001. Statistical analysis and generation of plots was performed using Prism v.7 software (GraphPad).

### Contact for reagent and resource sharing

Constructs and zebrafish transgenic lines available upon request to Maura McGrail. RNA-Seq data submission at GEO is in progress.

## Acknowledgements

The authors thank Marc Ekker, U of Ottawa for the *dat:eGFP* and *dat:mitoRFP* lines; Ed Boyden, Massachusetts Institute of Technology for the Higashijima lab *elavl3:Gal4* line; Diana Mitchell, University of Idaho for the 4C4 hybridoma supernatant; Raquel Espin, Iowa State University for the NFκΒ inhibitor CAPE; and Diane Slusarski, Rochester Institute of Technology, for use of the ZebraBox monitoring system and ZebraLab software.

## Data availability

RNASeq submission to GEO is in progress.

## Author contributions statement

SS: conceptualization, validation, formal analysis, investigation, writing – original draft preparation, writing – review and editing, visualization; FL: conceptualization, methodology, validation, formal analysis, investigation, writing – original draft preparation, writing – review and editing, visualization; AK: funding acquisition, writing – review and editing; MM: conceptualization, methodology, resources, writing – original draft preparation, writing – review and editing, visualization, supervision, project administration, funding acquisition.

